# Erythrocyte Rheology under Anesthesia: Insights from Glycated and Non-glycated Red Blood Cells

**DOI:** 10.1101/2025.09.15.676294

**Authors:** Marcus V. Batista Da Silva, Horacio V. Castellini, Nicolás A. Alet, Bibiana D. Riquelme, Analía I. Alet

**Affiliations:** Facultad Cs. Bioquímicas y Farmacéuticas, Universidad Nacional de Rosario, Suipacha 535, 2000 Rosario, Argentina; Facultad Cs. Exactas Ingeniería y Agrimensura, Universidad Nacional de Rosario, Pellegrini 250, 2000 Rosario, Argentina; Facultad Cs. Médicas, Universidad Nacional de Rosario, Santa Fe 3100, 2000 Rosario, Hospital Provincial del Centenario. Urquiza 3101, 2000 Rosario, Argentina; Grupo de Física Biomédica, IFIR (CONICET-UNR), Bv. 27 de febrero 210 bis, 2000 Rosario, Argentina; Consejo de Investigaciones de la UNR (CIUNR)

**Keywords:** Erythrocyte viscoelasticity, Erythrocyte glycation, Hemorheology, Propofol, Remifentanil, Vecuronium

## Abstract

This study investigated the effects of general anesthesia (propofol, vecuronium, remifentanil, and their combinations) on the aggregation and viscoelasticity of human erythrocytes, in both normal and *in vitro* glycated samples (simulating hyperglycemia as occurs in diabetes). Our results demonstrate that these anesthetics increase erythrocyte aggregation. Propofol and its combinations show a synergistic effect, forming larger aggregates. Analysis of the erythrocyte viscoelasticity revealed that propofol alone increased the elastic modulus, while the propofol+remifentanil+vecuronium combination decreased the storage modulus, suggesting complex interactions with the cytoskeleton and lipid bilayer. In glycated erythrocytes, the same anesthetic combinations do not significantly affect viscoelastic parameters. The membrane viscosity values of glycated erythrocytes were closer to those of the control. These findings highlight that these drugs affect hemorheologic parameters differently in non-glycated and glycated erythrocytes. These results provide valuable insights for understanding potential microvascular complications in diabetic patients during and after surgical procedures. We suggest expanding the study to the molecular level for a more comprehensive understanding of the chemical interactions between the drugs used during general anesthesia and the erythrocyte membrane.

## 1. INTRODUCTION

Diabetes mellitus encompasses a range of metabolic disorders characterized by hyperglycemia, resulting from defects in insulin secretion, insulin action, or both^1^. Persistent hyperglycemia leads to tissue damage and subsequent secondary complications^2^. According to the International Diabetes Federation, 9.3% of the adult population aged 20-79 had diabetes in 2019, with projections indicating an increase to 10.2% by 2030 and 10.9% by 2045, affecting approximately 700.2 million adults^3^. In individuals with diabetes, pathological changes such as alterations in erythrocyte surface charge, increased blood viscosity, heightened membrane stiffness, and enhanced formation of *rouleaux* and globular erythrocyte aggregates (clusters) can significantly impact hemorheological parameters and induce changes in microcirculation^4–10^. These alterations complicate the study of blood samples from diabetic patients, further complicated by comorbidities such as hypertension, obesity, and hyperlipidemia. Researchers have developed *in vitro* protocols to study the glycation process with greater control and accuracy to address these challenges^11^.

Whole blood and red blood cells (RBCs) are characterized as viscoelastic materials, displaying viscoelastic properties when subjected to oscillatory shear stresses. The principles of complex viscoelasticity describe this behaviour^12,13^. The dynamic nature of blood is significant because *in vivo* flow is pulsatile and non-stationary, influenced by the cardiac cycle and continuous changes in vessel size and flow conditions^14^. At rest, RBCs form aggregates resembling “coin stacks” known as “*rouleaux*.” Initiating blood flow requires an effort to disaggregate these erythrocytes, as blood initially behaves like a material with high viscosity. As erythrocytes disaggregate and flow begins, viscosity decreases^15,16^. In vessels, once laminar flow is established, erythrocytes align along streamlines and accumulate near the vessel axis. Plasma acts as a liquid sheath, further reducing viscosity, a phenomenon known as the Fahraeus-Lindquist effect^17^.

*In vitro* studies of blood samples from donors with type 2 diabetes mellitus (T2DM) treated with propofol have revealed significant hemorheological alterations compared to healthy individuals. Changes were observed in the viscoelastic properties of erythrocytes and in erythrocyte aggregation. These modifications are attributed to changes in surface electrical charge and, to a lesser extent, in the lipid bilayer^7,18^.

In surgical procedures requiring anesthetic agents, microvascular flow can be altered due to the effects of these drugs on the systemic cardiovascular system, impacting microcirculation. These changes often result from hemorheological disturbances such as increased platelet aggregation, viscosity and RBC deformability changes, elevated coagulation factors, and reduced fibrinolysis^18–20^. Anesthetics used in surgery can be administered either by inhalation or intravenously. Among intravenous anesthetics, the combination of propofol, remifentanil, and vecuronium is commonly employed^21^.

Given that hemorheological characteristics can differ between healthy individuals and those with pathological conditions, and that pathological changes in hemorheology and hemostasis may be detectable before and during surgery, it is crucial to implement preventive measures before surgical intervention to optimize patient outcomes. This study aims to explore the hemocompatibility of the anesthetics propofol (P), remifentanil (R), and vecuronium (V), both individually (P, R, V) and in combination (PR, PV, RV, and PRV), through detailed *in vitro* analysis. The objective is to understand how these agents, commonly used in general anesthesia, can alter erythrocyte morphology, viscoelasticity, aggregation, and deformability at steady-state concentrations, using human erythrocytes from healthy donors (control, C) as well as *in vitro* glycated erythrocytes (g) to model diabetes-related changes.

## 2. RESULTS AND DISCUSSION

### 2.1 Effects of Anesthetics Drugs on RBCs

#### 2.1.1 Digital Analysis of Microscopic Images

In order to characterize the effects of these drugs used during general anesthesia on the membrane of non-glycated RBCs, incubations were performed with different anesthetic drugs, individually and in combination, as described in Table 9 of the Materials and Methods section. Figure 1 shows representative images for each incubation condition, where an increase in *rouleaux* formation can be observed following exposure to these agents.

**Figure 1.**
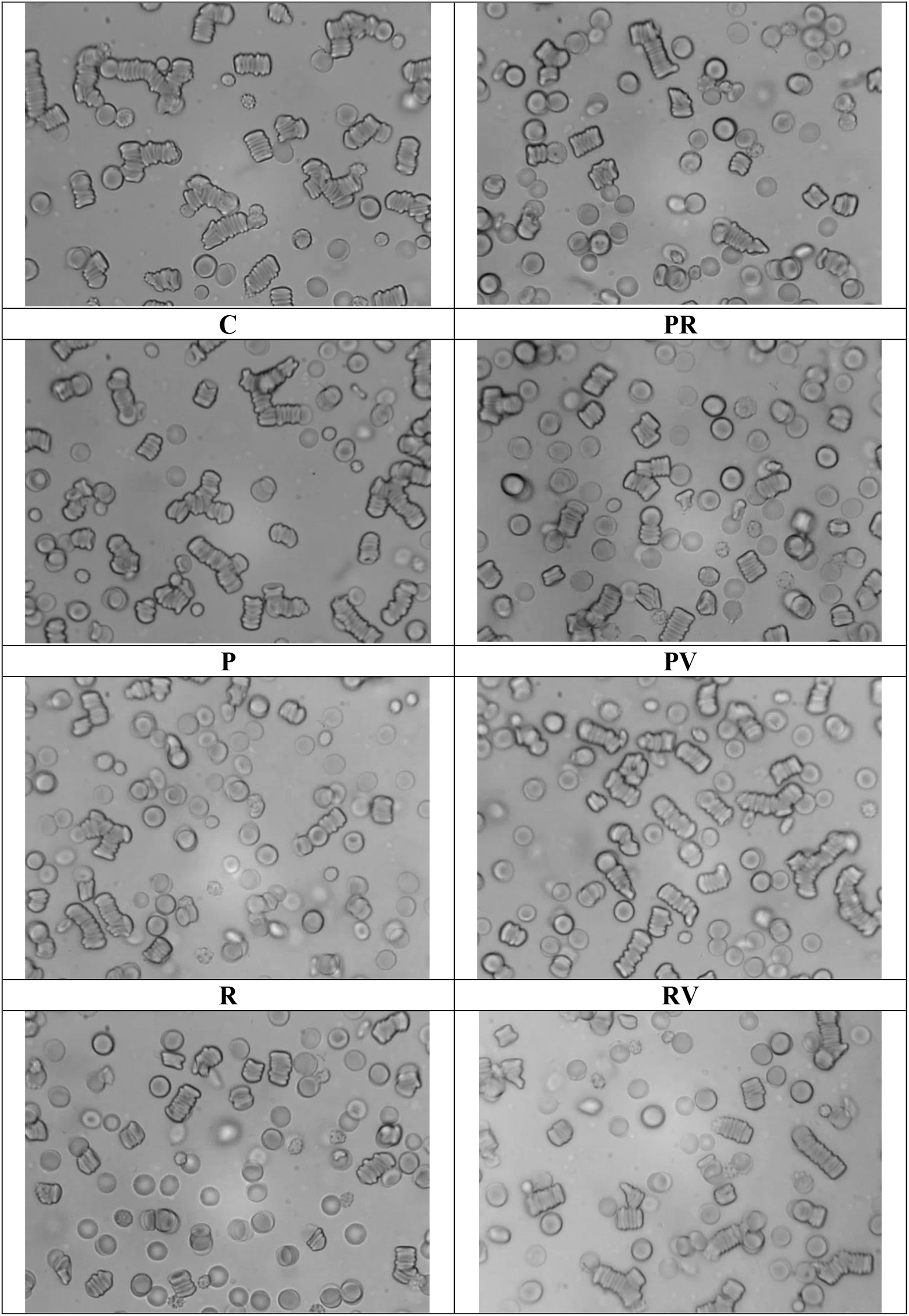
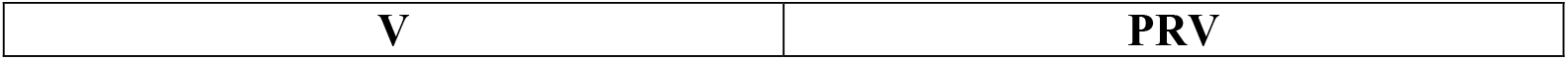
Representative microscopic images of RBCs samples (non-glycated) treated with anesthetics.

Table 1 shows the percentages for each type of erythrocyte aggregate formation, using the classification criteria based on the number of cells composing the aggregates^22,23^.

**Table 1.** Percentage of RBC (non-glycated) aggregate categories treated with anesthetics (Mean ± standard deviation).

| Samples | % I.C. | % 2, 3 or 4 RBCs | % 5 or more RBCs | % AMAS |
| --- | --- | --- | --- | --- |
| C | 28 ± 3 | 49 ± 3 | 20 ± 3 | 3 ± 2 |
| P | 26 ± 4 | 39 ± 5 | 30 ± 5* | 5 ± 2 |
| R | 28 ± 5 | 48 ± 2 | 20 ± 6 | 4 ± 1 |
| V | 27 ± 6 | 43 ± 3 | 27 ± 4* | 3 ± 1 |
| PR | 27 ± 2 | 42 ± 3* | 24 ± 6 | 7 ± 2 |
| PV | 26 ± 2 | 41 ± 5* | 27 ± 4* | 6 ± 2 |
| RV | 24 ± 5 | 45 ± 6 | 26 ± 5 | 5 ± 2 |
| PRV | 24 ± 3 | 42 ± 2** | 26 ± 3* | 8 ± 1** |

\* $p < 0.05$ ; \*\* $p < 0.01$ ; \*\*\* $p < 0.001$ compared to C

After incubation with these agents, longer *rouleaux* formations were observed. For example, the C shows a predominance of aggregates of up to 4 stacked RBCs (49%) and a very scarce presence of larger aggregates (AMAS). At the same time, all samples treated with drugs used during general anesthesia exhibit lower percentages of aggregates of up to 4 cells. This decrease is significant for the PR, PV, and PRV combinations. When analyzing the formation of aggregates with five or more RBCs, a significant increase is observed in the P and V samples compared to the control (30% and 27%, respectively). This behaviour is also seen in mixtures containing both P and V. For instance, the PV and PRV samples show significant differences in the percentage of aggregates with five or more RBCs (27% and 26%, respectively) compared to the control. These results suggest that these drugs used during general anesthesia may promote RBC aggregation.

Figure 2 shows the CCA values^24^, where it is observed that the samples treated with P and V separately, as well as the PV and PRV mixtures, exhibit significantly higher aggregation than the control, approaching a value of 1 (complete aggregation). This finding is consistent with the results presented in Table 1 and the images in Figure 1.

**Figure 2.**
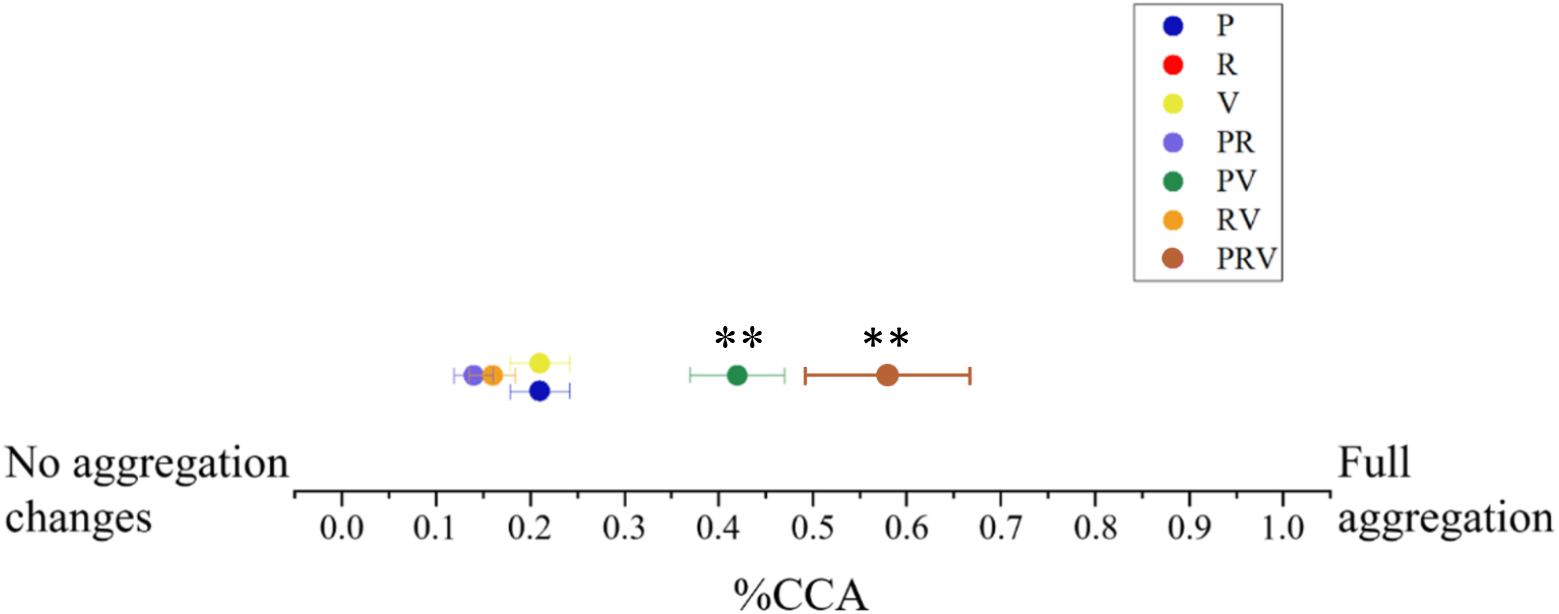
Isolated Cells Coefficient (CCA) parameter for non-glycated RBCs treated with different anesthetics compared to Control. *p<0.05; **p<0.01; ***p<0.001.

#### 2.1.2 Aggregation Kinetics Parameters

Table 2 presents the aggregation kinetics parameters obtained with the optical chip aggregometer. Amp_600_ does not show significant differences for the samples treated with drugs used during anesthesia. However, the half-time (t_1/2_) shows a significant decrease for the PV, RV, and PRV samples compared to the control, indicating that the agents treatment could accelerate RBC aggregation. Regarding the Aggregation Index (AI), a significant decrease is observed for R and a significant increase for RV and PRV compared to C.

**Table 2.** Aggregation kinetics parameters for non-glycated RBCs treated with anesthetics. (Mean ± standard deviation).

| Samples | Amp<sub>600</sub><br>a. u. | $t_{1/2}$<br>(s) | AI<br>a. u. |
| --- | --- | --- | --- |
| C | 82 ± 5 | 77 ± 3 | 0.74 ± 0.05 |
| P | 83 ± 4 | 66 ± 4 | 0.76 ± 0.03 |
| R | 80 ± 2 | 75 ± 5 | 0.71 ± 0.02* |
| V | 83 ± 4 | 65 ± 4 | 0.75 ± 0.03 |
| PR | 80 ± 4 | 71 ± 4 | 0.72 ± 0.02 |
| PV | 79 ± 6 | 69 ± 7* | 0.72 ± 0.04 |
| RV | $87 \pm 7$ | $61 \pm 9^{**}$ | $0.80 \pm 0.07^*$ |
| PRV | $88 \pm 4$ | $50 \pm 2^{***}$ | $0.81 \pm 0.05^*$ |

*\*p<0.05; \*\*p<0.01; \*\*\*p<0.001 compared to C*

#### 2.1.3 Stationary and Dynamic Viscoelastic Parameters

Table 3 presents the stationary viscoelastic parameters of control RBCs and those treated with anesthetics obtained with the Erythrocyte Rheometer. When analyzing the membrane surface viscosity (η) and elastic modulus (µ), a significant increase is observed in the P sample compared to C. However, the deformability index (DI) shows a significant decrease for the sample treated with P compared to C. These results suggest that the membrane could be altered by incubation with P, which could lead to microcirculation disturbances.

**Table 3.** Stationary viscoelastic parameters for non-glycated RBC samples treated with anesthetics. (Mean ± standard deviation).

| Samples | $\eta$<br>$10^{-7}\text{N.s/m}$ | $\mu$<br>$10^{-6}\text{N/m}$ | DI |
| --- | --- | --- | --- |
| **C** | $2.1 \pm 0.5$ | $4.9 \pm 0.2$ | $0.64 \pm 0.02$ |
| **P** | $2.3 \pm 0.7^*$ | $5.06 \pm 0.08^{***}$ | $0.62 \pm 0.01^{**}$ |
| **R** | $2.1 \pm 0.7$ | $4.8 \pm 0.2$ | $0.64 \pm 0.01$ |
| **V** | $2.0 \pm 0.4$ | $4.9 \pm 0.1$ | $0.63 \pm 0.02$ |
| **PR** | $2.1 \pm 0.2$ | $4.9 \pm 0.2$ | $0.63 \pm 0.01^*$ |
| **PV** | $2.2 \pm 0.3^*$ | $4.9 \pm 0.1$ | $0.63 \pm 0.01^{***}$ |
| **RV** | $1.9 \pm 0.3$ | $4.8 \pm 0.1$ | $0.69 \pm 0.01^{***}$ |
| **PRV** | $1.9 \pm 0.3$ | $4.80 \pm 0.02^{***}$ | $0.62 \pm 0.01^{***}$ |

*\*p<0.05; \*\*p<0.01; \*\*\*p<0.001 compared to C.*

It can be observed that the samples treated with V and R do not show significant differences in any of the stationary viscoelastic parameters compared to C. However, when analyzing the PR, PV, and PRV combinations, a significant decrease is observed in the DI, suggesting that these changes might be associated with using P. Additionally, the PRV samples show a significant decrease in the RBC membrane elastic modulus µ, revealing a reduction in erythrocyte membrane stiffness due to the action of combined agents, yielding values close to those of C.

Oscillatory measurements allow for a more detailed description of the behaviour of RBCs under conditions similar to blood circulation. Using the heart rate as a reference, at 60 beats per minute (equivalent to a frequency of 1 Hz), and considering that RBCs can be classified as viscoelastic materials, we can analyze the phase shift between the applied shear stress and the erythrocyte response. Figure 3 shows the phase shifts (δ) between the shear stress and the erythrocyte response at a frequency of 1 Hz. The PRV sample has the most considerable phase shift, but the difference is not significant (p > 0.05).

**Figure 3.**
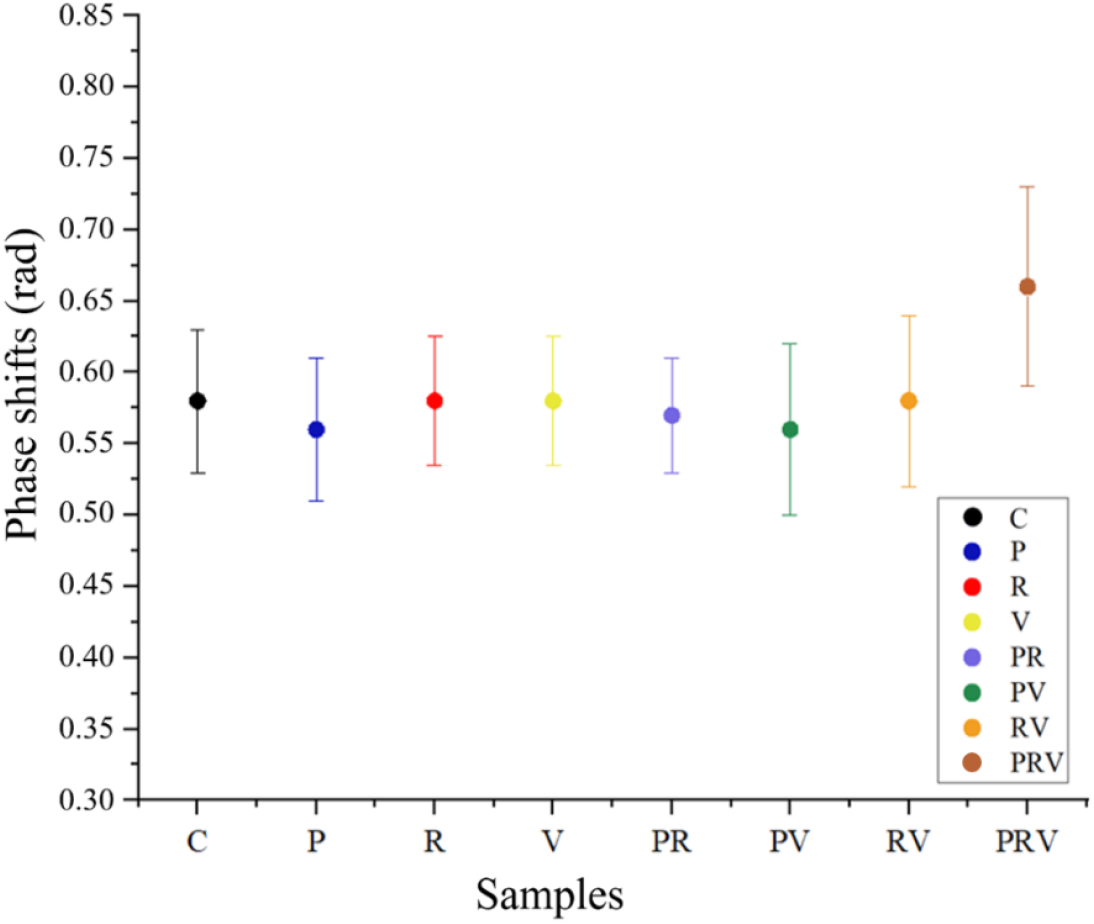
Phase shift δ between shear stress and deformation at an oscillatory frequency of 1.0Hz. p>0.05 in all cases.

Table 4 presents the dynamic viscoelastic parameters and the elastic modulus of the RBC membrane. It can be observed that the storage modulus (G’) shows a significant decrease for the PRV-treated samples compared to the control (C). The decrease in the storage modulus (G’) could be related to an alteration in the cytoskeleton. This is consistent with the loss modulus (G”) changes, where a significant increase is observed for the same PRV sample, indicating greater energy dissipation than C. These results suggest interactions between the combined anesthetics and the lipid bilayer may occur.

**Table 4.** Dynamic viscoelastic parameters for non-glycated RBC samples treated with anesthetics (oscillatory frequency of 1.0 Hz). (Mean ± standard deviation).

| Samples | $G'$<br>$10^{-6}\text{N/m}$ | $G''$<br>$10^{-6}\text{N/m}$ | $\eta'$<br>$10^{-7}\text{N.s/m}$ | $\eta''$<br>$10^{-7}\text{N.s/m}$ |
| --- | --- | --- | --- | --- |
| **C** | $4.11 \pm 0.09$ | $2.6 \pm 0.2$ | $4.2 \pm 0.3$ | $6.7 \pm 0.4$ |
| **P** | $4.08 \pm 0.05$ | $2.64 \pm 0.05$ | $4.21 \pm 0.02$ | $6.6 \pm 0.4$ |
| **R** | $4.12 \pm 0.05$ | $2.6 \pm 0.3$ | $4.3 \pm 0.2$ | $6.6 \pm 0.7$ |
| **V** | $4.1 \pm 0.1$ | $2.6 \pm 0.1$ | $4.2 \pm 0.4$ | $6.4 \pm 0.2$ |
| **PR** | $4.2 \pm 0.2$ | $2.7 \pm 0.2$ | $4.3 \pm 0.2$ | $6.7 \pm 0.2$ |
| **PV** | $4.1 \pm 0.1$ | $2.7 \pm 0.1$ | $4.3 \pm 0.3$ | $6.6 \pm 0.3$ |
| **RV** | $4.06 \pm 0.05$ | $2.69 \pm 0.02$ | $4.29 \pm 0.08$ | $6.54 \pm 0.04$ |
| **PRV** | $3.74 \pm 0.08^{***}$ | $2.93 \pm 0.02^{***}$ | $4.7 \pm 0.2^{**}$ | $5.96 \pm 0.08^*$ |

\* $p < 0.05$ ; \*\* $p < 0.01$ ; \*\*\* $p < 0.001$ compared to C.

### 2.2 Effect of Incubation with Anesthetics on Glycated RBCs

To characterize the effects of drugs on the membrane of glycated RBCs, incubations were carried out using the same anesthetic agents—individually and in combination—as described in Table 9 of the Materials and Methods section. The following hemorheological determinations examine the impact of these drugs on glycated erythrocytes, highlighting notable alterations in aggregation behaviour and viscoelastic properties.

#### 2.2.1 Digital Analysis of Microscopic Images

Figure 4 shows representative images of glycated samples treated with different anesthetics, alone or in combination. It can be observed that all samples subjected to the glycation process exhibit larger aggregates than those incubated only with anesthetics (Figure 1).

**Figure 4.**
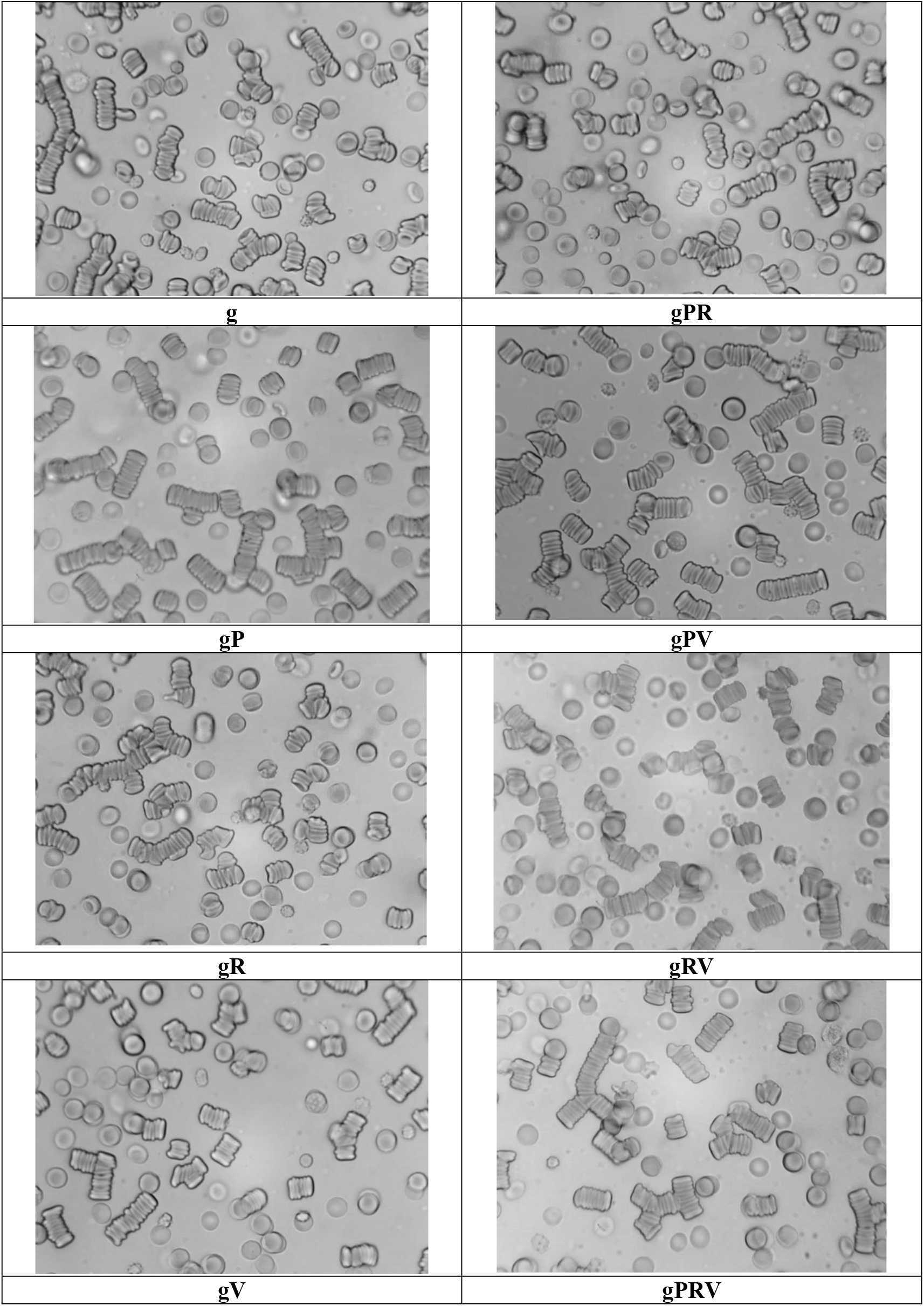
Representative microscopic images of glycated red blood cell samples treated with anesthetics.

Table 5 presents the percentage distribution of glycated RBC aggregates treated with anesthetics. It shows that for glycated RBCs (g), 74% of the aggregates correspond to the distribution of 2, 3, or 4 cells (32%) and five or more cells (42%).

**Table 5.** Percentage of aggregate categories of glycated RBCs treated with different anesthetics. (Mean ± standard deviation).

| Samples | % I.C. | % 2, 3 or 4 RBCs | % 5 or more RBCs | % AMAS |
| --- | --- | --- | --- | --- |
| **C** | 28 ± 3 | 49 ± 3 | 20 ± 3 | 3 ± 2 |
| **g** | 19 ± 2 | 32 ± 3 | 42 ± 5 | 7 ± 2 |
| **gP** | 15 ± 4 | 26 ± 4 | 47 ± 5 | *12 ± 4 |
| **gR** | 25 ± 4** | 27 ± 2* | 41 ± 4 | 7 ± 3 |
| **gV** | 20 ± 3 | 33 ± 5 | 44 ± 6 | 3 ± 1* |
| **gPR** | 22 ± 4 | 26 ± 4 | 43 ± 4 | 9 ± 3 |
| **gPV** | 19 ± 2 | 27 ± 2 | 45 ± 3 | 9 ± 2* |
| **gRV** | 18 ± 3 | 26 ± 2 | 45 ± 2 | 11 ± 2* |
| **gPRV** | 15 ± 2* | 23 ± 3** | 48 ± 6 | 14 ± 2** |

\* $p < 0.05$ ; \*\* $p < 0.01$ ; \*\*\* $p < 0.001$ compared to g.

In Table 5, it can be observed that all glycated samples treated with anesthetics exhibit larger aggregates compared to the control. Notably, the gR and gPRV samples show a significant decrease in the percentage of aggregates with up to 4 RBCs compared to the g sample. Additionally, the gP, gPV, gRV, and gPRV samples presented a significant increase in AMAS formation compared to g. At the same time, for the Vg treatment, this percentage decreased, reaching the value of the control (C). This would indicate that the anesthetics exhibit a synergistic effect with glucose, increasing erythrocyte aggregation, except for the incubation with V.

In the isolated cell coefficient for the different treatments (Figure 5), it can be seen that gPV and gPRV showed a significant increase in the %CCA value compared to g, indicating greater aggregation in these samples than those not treated with anesthetics.

**Figure 5.**
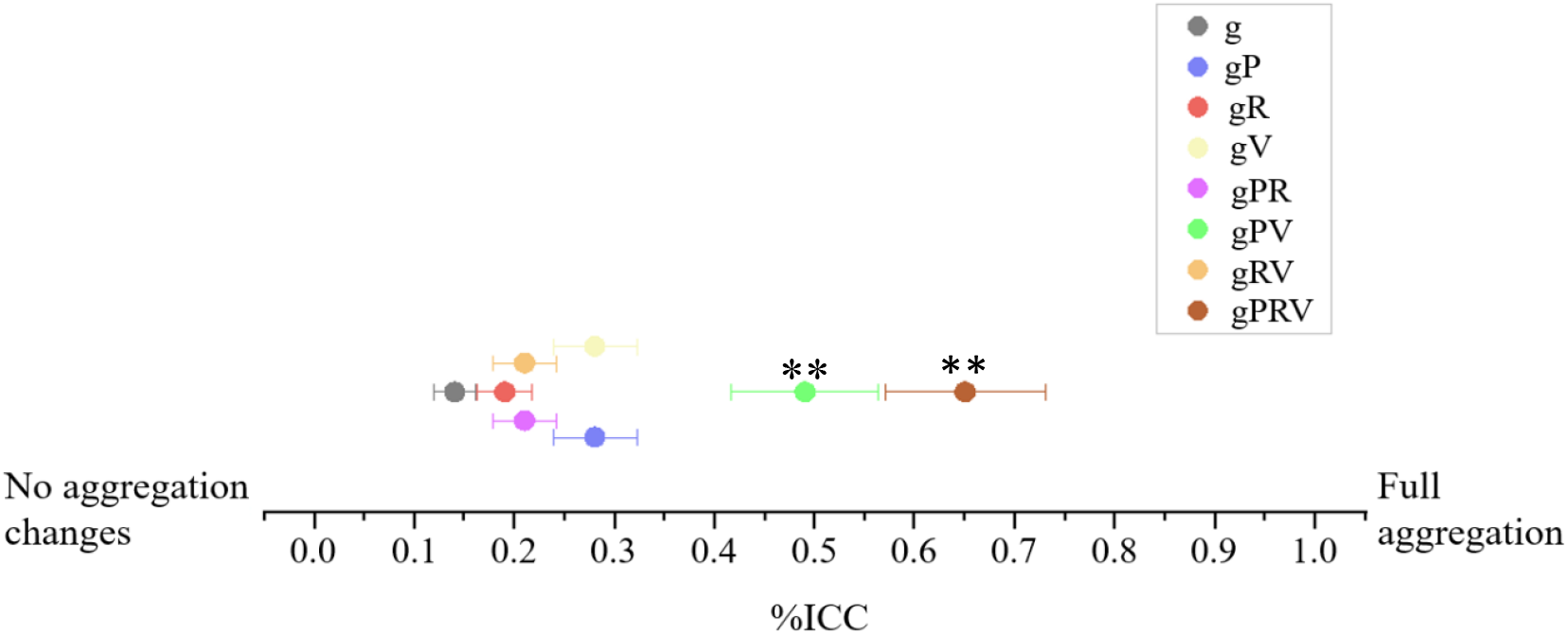
CCA parameter for glycated RBCs treated with anesthetics. *p<0.05; **p<0.01; *p<0.001 compared to g.

#### 2.2.2 Aggregation Kinetics Parameters

Table 6 presents the aggregation kinetics parameters, where it can be observed that Amp600 does not show significant differences for glycated samples treated with anesthetics. On the other hand, the half-time shows a significant decrease for the gPV and gPRV samples, indicating that both glucose and the anesthetics used may accelerate RBC aggregation. However, the gRV sample showed a significant increase in the half-time compared to g. It exceeded the C value, suggesting that this combination of anesthetics could slow down the aggregation rate of glycated RBCs.

**Table 6.** Aggregation kinetics parameters for glycated and non-glycated samples treated with anesthetics. (Mean ± standard deviation).

| Samples | Amp<sub>600</sub><br>a. u. | t<sub>1/2</sub><br>(s) | AI<br>a. u. |
| --- | --- | --- | --- |
| C | 82 ± 5 | 77 ± 3 | 0.74 ± 0.05 |
| g | 82 ± 6 | 69 ± 3 | 0.75 ± 0.04 |
| gP | 84 ± 7 | 62 ± 6 | 0.76 ± 0.07 |
| gR | 83 ± 5 | 73 ± 4 | 0.75 ± 0.08 |
| gV | 81 ± 2 | 63 ± 2 | 0.74 ± 0.02 |
| gPR | 86 ± 7 | 70 ± 7 | 0.77 ± 0.06 |
| gPV | 85 ± 8 | 52 ± 7* | 0.78 ± 0.08 |
| gRV | 88 ± 2 | 82 ± 5* | 0.79 ± 0.03 |
| **gPRV** | $89 \pm 4$ | $47 \pm 5^{***}$ | $0.81 \pm 0.04^*$ |

*\*p<0.05; \*\*p<0.01; \*\*\*p<0.001 compared to g*

#### 2.2.3 Stationary and Dynamic Viscoelastic Parameters

Table 7 shows the stationary viscoelastic parameters, and it can be seen that gP exhibited a significant increase in the membrane surface viscosity compared to the g sample. On the other hand, the gPR, gRV, and gPRV samples showed a significant decrease for the same parameter. Notably, an increase in membrane viscosity was observed with the P treatment. However, when combined with R and V, an antagonistic effect was seen, significantly reducing this parameter. It can also be observed that the elastic modulus of the membrane significantly decreased for the samples treated with gR, gV, gPV, gRV, and gPRV compared to g. In line with this, the ID showed a significant decrease for the gPRV sample. On the other hand, the gR, gV, gPV, and gRV samples showed a significant increase in DI.

**Table 7.** Stationary viscoelastic parameters for glycated samples treated with anesthetics. (Mean ± standard deviation).

| **Samples** | **$\eta$**<br>**$10^{-7}\text{N.s/m}$** | **$\mu$**<br>**$10^{-6}\text{N/m}$** | **DI** |
| --- | --- | --- | --- |
| **C** | $2.1 \pm 0.5$ | $4.9 \pm 0.2$ | $0.64 \pm 0.02$ |
| **g** | $2.4 \pm 0.6$ | $5.11 \pm 0.04$ | $0.62 \pm 0.01$ |
| **gP** | $2.9 \pm 0.2^*$ | $5.13 \pm 0.03$ | $0.63 \pm 0.02$ |
| **gR** | $2.1 \pm 0.3$ | $5.03 \pm 0.08^{***}$ | $0.64 \pm 0.03^*$ |
| **gV** | $2.1 \pm 0.4$ | $5.00 \pm 0.05^{**}$ | $0.64 \pm 0.02^{***}$ |
| **gPR** | $1.9 \pm 0.3^*$ | $5.09 \pm 0.01$ | $0.63 \pm 0.01$ |
| **gPV** | $2.5 \pm 0.5$ | $5.06 \pm 0.06^{***}$ | $0.64 \pm 0.02^{**}$ |
| **gRV** | $2.0 \pm 0.2^{***}$ | $5.05 \pm 0.09^*$ | $0.65 \pm 0.02^*$ |
| **gPRV** | $1.9 \pm 0.4^{***}$ | $4.85 \pm 0.01^{***}$ | $0.61 \pm 0.01^{***}$ |

*\*p<0.05; \*\*p<0.01; \*\*\*p<0.001 compared to g.*

Figure 6 presents the phase shifts δ between the applied shear stress and the erythrocyte response at a frequency of 1 Hz for glycated RBCs incubated with anesthetics, showing no significant differences.

**Figure 6.**
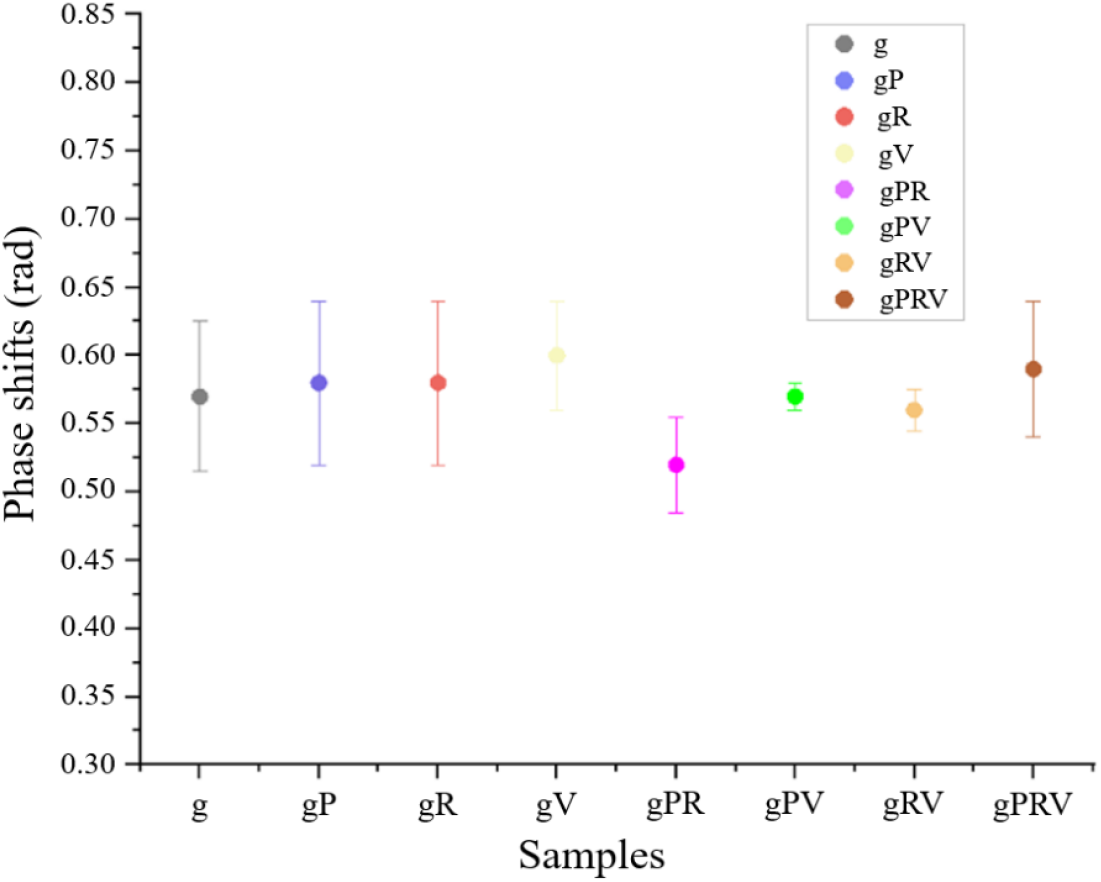
Phase difference δ between stress and strain for oscillatory frequencies of 1.0 Hz. p>0.05 in all cases.

Table 8 presents the dynamic viscoelastic parameters. The storage modulus G’ shows a significant decrease for the samples treated with gR, gV, gPV, and gPRV compared to the g sample, with these values being close to those of the C. The dynamic loss modulus G” shows a significant decrease for the gP, gPR, gRV, and gPRV samples compared to g, suggesting that it takes longer to dissipate energy. On the other hand, it can be observed that the real component of complex viscosity η’ shows significant differences for the gP, gR, gPR, and gPRV samples. In contrast, the imaginary component η” indicates a significant decrease for the gV and gPRV samples. This is consistent with what was observed for G’ and G”. These results show alterations compared to the g sample, with the dynamic viscoelastic parameters being similar to the values of the C sample, suggesting that treatment with anesthetics in glycated RBC could reverse the effects of glycation.

**Table 8.** Dynamic viscoelastic parameters for glycated RBC samples treated with anesthetics at an oscillation frequency of 1.0 Hz. (Mean ± standard deviation).

| Samples | $G'$<br>$10^{-6}\text{N/m}$ | $G''$<br>$10^{-6}\text{N/m}$ | $\eta'$<br>$10^{-7}\text{N.s/m}$ | $\eta''$<br>$10^{-7}\text{N.s/m}$ |
| --- | --- | --- | --- | --- |
| **C** | 4.11 ± 0.09 | 2.6 ± 0.2 | 4.2 ± 0.3 | 6.7 ± 0.4 |
| **g** | 4.32 ± 0.08 | 2.7 ± 0.2 | 4.3 ± 0.1 | 6.83 ± 0.06 |
| **gP** | 4.4 ± 0.3 | 2.55 ± 0.08* | 4.06 ± 0.07** | 6.9 ± 0.5 |
| **gR** | 4.28 ± 0.02* | 2.6 ± 0.2 | 4.16 ± 0.06* | 6.8 ± 0.2 |
| **gV** | 4.18 ± 0.03** | 2.7 ± 0.1 | 4.3 ± 0.4 | 6.65 ± 0.04** |
| **gPR** | 4.4 ± 0.3 | 2.52 ± 0.06* | 4.0 ± 0.1* | 7.0 ± 0.2 |
| **gPV** | 4.26 ± 0.04* | 2.7 ± 0.3 | 4.3 ± 0.3 | 6.78 ± 0.04 |
| **gRV** | 4.27 ± 0.08 | 2.67 ± 0.04* | 4.3 ± 0.3 | 6.81 ± 0.3 |
| **gPRV** | 4.18 ± 0.03*** | 2.4 ± 0.02** | 3.8 ± 0.3** | 6.69 ± 0.03*** |

\* $p < 0.05$ ; \*\* $p < 0.01$ ; \*\*\* $p < 0.001$ compared to *g*.

**Table 9.** Anesthetic Incubations with Glycated and Non-Glycated RBCs.

| RBCs<br>(C) | RBCs glycated<br>(g) | Propofol<br>(P) | Remifentanil<br>(R) | Vecuronium<br>(V) | PBSA | Plasma | Name |
| --- | --- | --- | --- | --- | --- | --- | --- |
| 200µL |  |  |  |  | 500µL | 300µL | C |
|  | 200µL |  |  |  | 500µL | 300µL | g |
| 200µL |  | 500µL |  |  |  | 300µL | P |
|  | 200µL | 500µL |  |  |  | 300µL | gP |
| 200µL |  |  | 500µL |  |  | 300µL | R |
|  | 200µL |  | 500µL |  |  | 300µL | gR |
| 200µL |  |  |  | 500µL |  | 300µL | V |
|  | 200µL |  |  | 500µL |  | 300µL | gV |
| 200µL |  | 500µL | *1.2µL |  |  | 300µL | PR |
|  | 200µL | 500µL | *1.2µL |  |  | 300µL | gPR |
| 200µL |  | 500µL |  | †9.0µL |  | 300µL | PV |
|  | 200µL | 500µL |  | †9.0µL |  | 300µL | gPV |
| 200µL |  |  | *1.2µL | 500µL |  | 300µL | RV |
|  | 200µL |  | *1.2µL | 500µL |  | 300µL | gRV |
| 200µL |  | 500µL | *1.2µL | †9.0µL |  | 300µL | PRV |
|  | 200µL | 500µL | *1.2µL | †9.0µL |  | 300µL | gPRV |

\* Concentrated at 5 µg/mL, followed by an additional aliquot after 15 minutes of incubation.
† Concentrated at 10 µg/mL

## 3. CONCLUSION

### 3.1 On the effects of anesthetics on human RBCs

The results obtained in this study demonstrate that all human RBC samples treated *in vitro* with the anesthetics P, V, PR, PV, and PRV exhibit increased erythrocyte aggregation. This is evidenced by a decrease in the proportion of small aggregates (up to 4 RBCs) and a corresponding increase in larger aggregates. Particularly, it is observed that in the P, PR, PV, and PRV samples, the aggregation is increased, suggesting that this alteration is due to P and a synergistic effect when combined with the other drugs. Another finding corroborating the previous observations is the CCA parameter, which shows a significant increase for the PV and PRV samples. The synergistic effect of P was also confirmed through the study with the optical chip aggregometer, where the t_1/2_ was significantly lower for PV and PRV, indicating that this combination of anesthetics would accelerate the erythrocyte aggregation process.

The RBCs treated solely with P exhibited an increase in the elastic modulus of the erythrocyte membrane. However, the RBC samples treated with the PRV combination showed a decrease in the storage modulus (G’) and an increase in the dynamic loss (G”), suggesting interactions between the combined anesthetics with the cytoskeleton and the lipid bilayer.

### 3.2 On the effects of anesthetics on glycated RBCs

The digital image analysis of the gPV, gRV, and gPRV samples showed a significant increase in aggregation, particularly in forming AMAS. It was also observed that the gPV and gPRV samples showed a significant increase in the CCA value, indicating that, in addition to forming large aggregates, they had fewer isolated RBCs after incubation with the anesthetics. The lowest t_1/2_ values were obtained for the gPV and gPRV samples, indicating that this combination accelerated the aggregation of the glycated RBCs. A synergistic action of glycation and anesthetics could explain the increase in size and speed of aggregate formation.

The gP sample showed increased membrane surface viscosity, but the gPR, gRV, and gPRV samples showed a significant decrease compared to the g sample. The values obtained for these samples are closer to the C, indicating that while the treatment of glycated RBCs with the combined anesthetics may decrease the membrane surface viscosity, this decrease brings it closer to the control value. Furthermore, a significant decrease in the elastic modulus of the glycated RBC membrane treated with anesthetics was observed for the following samples: gR, gV, gPV, gRV, and gPRV, which also approached the control values. These results indicate that the glycated RBCs did not adversely affect the viscoelastic parameters due to incubation with the anesthetics. On the contrary, they moved closer to the values found for the control RBCs.

### 3.3 General Conclusions and Future Perspectives

The results show that the hemorheological activity of anesthetic agents affects erythrocyte aggregation and viscoelasticity in different ways, both in glycated and non-glycated RBCs.

These findings provide valuable insights not only for a better understanding of the hemorheological effects of anesthetics but also for the preventive assessment of potential complications during surgical procedures and postoperative microvascular dysfunction in diabetic patients.

For a more comprehensive understanding of the interaction between anesthetic agents and RBCs (glycated or non-glycated), it would be interesting to expand the study from a chemical perspective by conducting experiments at the molecular level. This approach could broaden the proposed model and elucidate how anesthetics interact with erythrocyte membrane proteins, particularly when glycated.

## 4. MATERIALS AND METHODS

To evaluate the effects of the anesthetics on hemorheological properties, a detailed experimental workflow was employed, as schematically illustrated in Figure 7. The procedure consisted of the *in vitro* glycation of erythrocytes to model hyperglycemia, followed by the incubation of the samples with the anesthetic agents. Finally, hemorheological parameters were determined for both the glycated and non-glycated groups.

**Figure 7.**
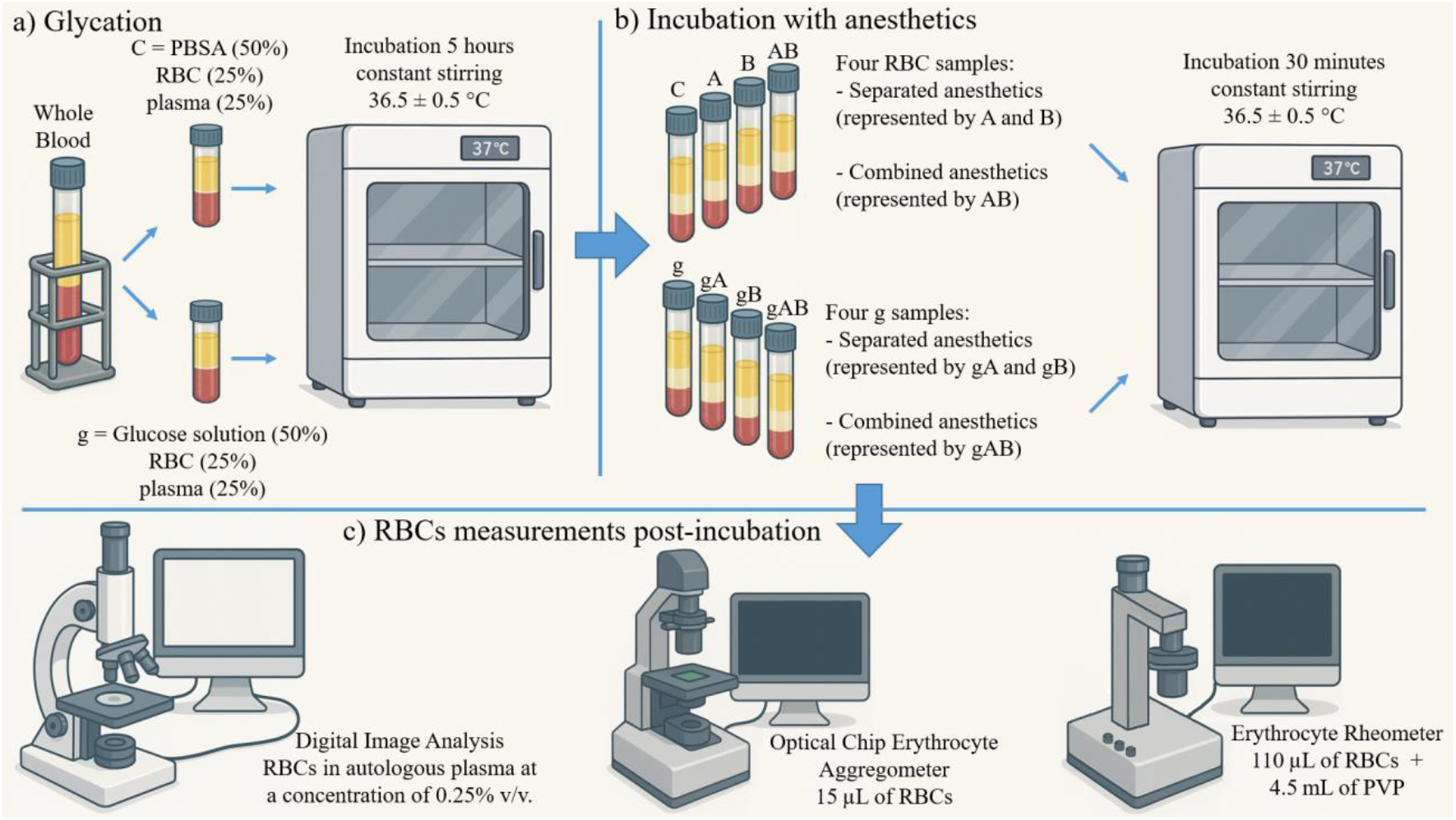
Schematic diagram of the experimental workflow. The diagram outlines the key steps of the study, from sample preparation to data acquisition. (a) In vitro glycation process to model the effects of hyperglycemia on red blood cells (RBCs). (b) Incubation of the glycated and non-glycated RBCs with individual and combined anesthetic agents. (c) Post-incubation analysis and data acquisition of hemorheological parameters to assess the effects of the treatments.

### 4.1 Biological Samples

Human blood samples from healthy donors (8 hs fasting, non-smoking, both sexes, aged between 20 and 35 years), n=15, who had not taken any medications in the last 7 days and had no known pre-existing or significant health problems, were collected by venipuncture in sterile vials containing EDTA as anticoagulant. Collection and processing of samples were performed within four hours of extraction time^25^.

The extraction and processing of the samples were conducted following the recommendations of the expert panel on hemorheology from the International Council for Standardization in Hematology, as well as the Bioethics regulations of the Facultad de Ciencias Bioquímicas y Farmacéuticas of the Universidad Nacional de Rosario (Res. No. 347/2013 and 425/2017), and all donors signed the informed consent.

Initially, the blood was centrifuged to separate and collect the plasma (1,500 g, 5 min, at room temperature), and the buffy coat was removed. After separating the plasma, the RBCs were washed twice with phosphate buffer solution (PBS; 137 mM NaCl, 1.4 mM KH_2_PO_4_, 10 mM Na_2_HPO_4_, 2.7 mM KCl, pH 7.4, 300 mOsM). Subsequently, the blood sample’s hematocrit was adjusted to 40% using autologous plasma.

### 4.2 Chemicals and Tools table

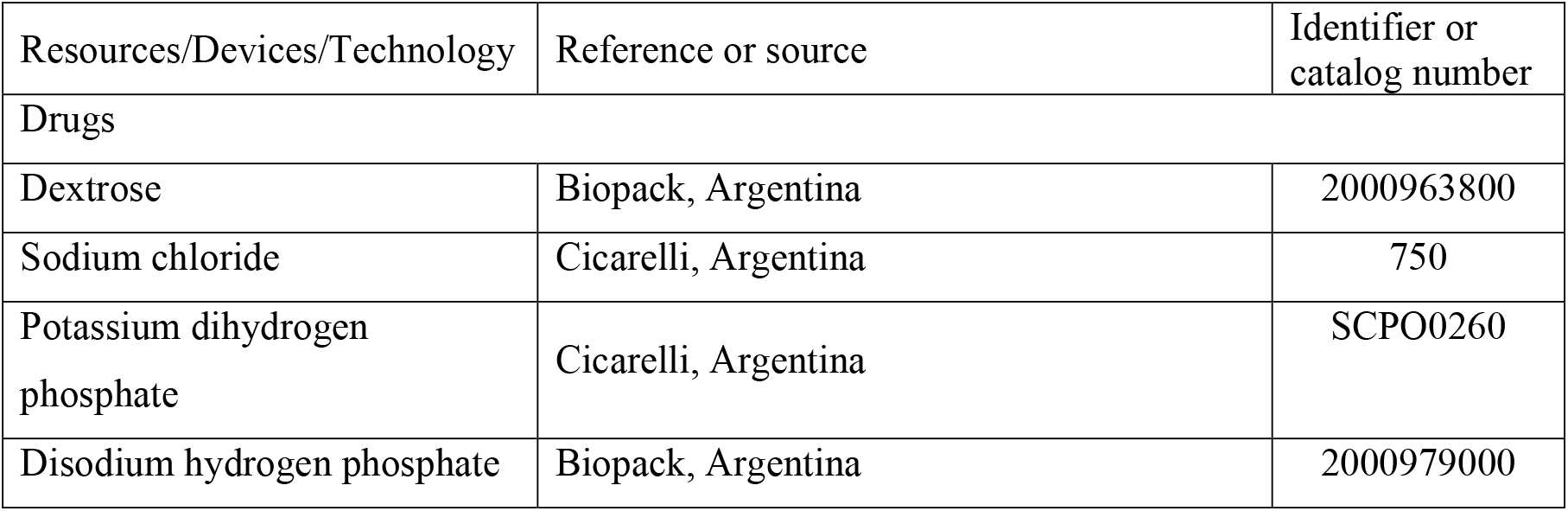
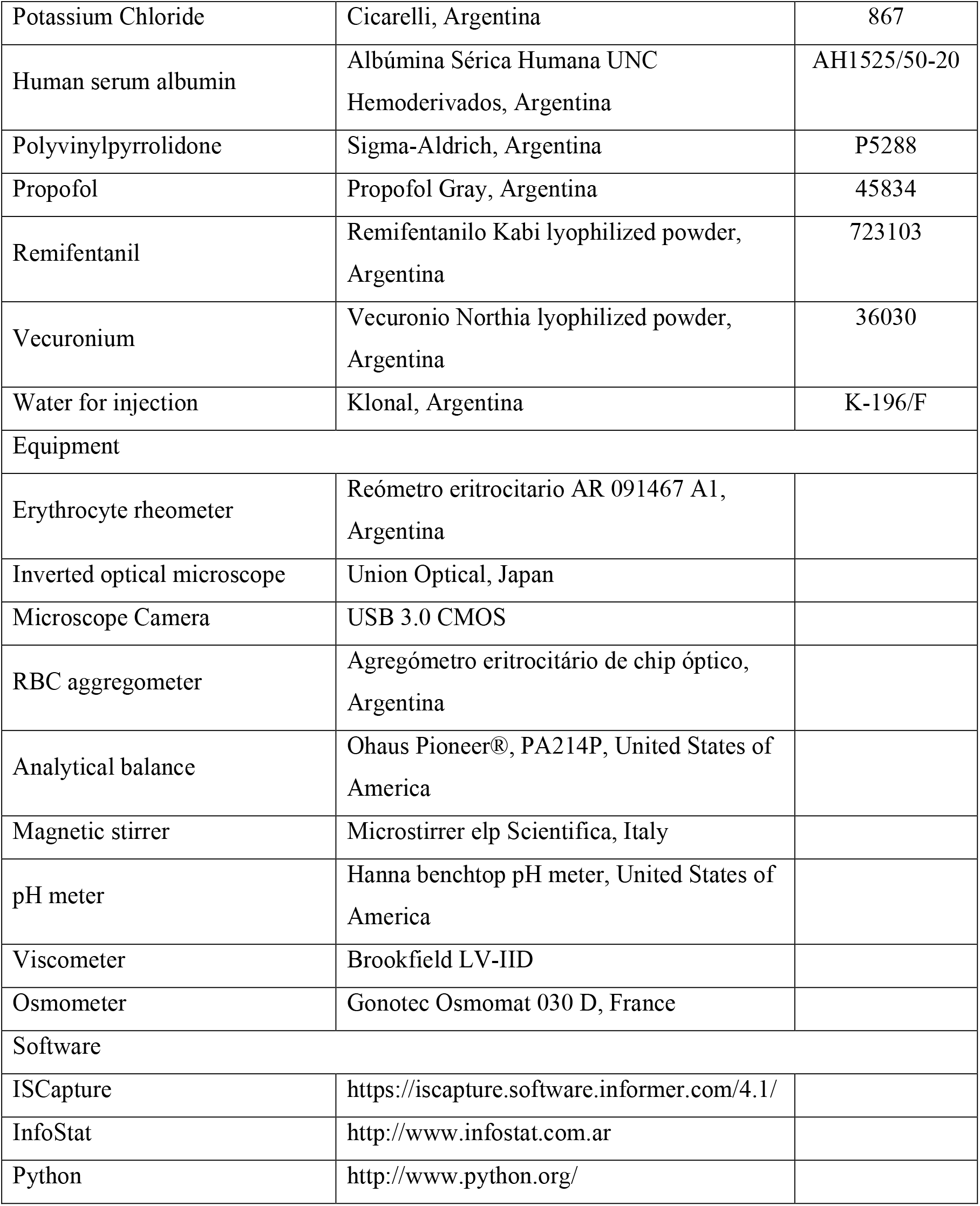

### 4.3 *In vitro* RBCs glycation

For glycation (g), a strict protocol was followed^11^. The incubation medium was prepared by dissolving 0.04 g of dextrose in 0.02 dL of PBSA (PBS supplemented with Human Serum Albumin 1% v/v). The samples were obtained by mixing equal volumes of washed RBCs and the incubation medium, resulting in a final glucose concentration of 1 g/dL and a hematocrit of 50%. Incubation was conducted for 5 hours at 36.5 °C with constant slow stirring. After this period, the RBCs were washed twice with PBS using gentle centrifugation (1,500 g, 5 min, at room temperature). The RBCs packages were reserved for further analysis. The control sample (C) consisted of RBCs incubated only with PBSA.

### 4.4 Anesthetic solutions for RBC incubation

These anesthetic concentrations in this study matched physiological levels used during anesthesia (steady-state concentrations) as described in the literature^26^. The following anesthetic solution were prepared:

- Propofol commercial suspension was diluted in PBSA to obtain a concentration of 8 μg/mL.
- Remifentanil was diluted in PBSA to obtain a concentration of 12 ng/mL (corresponding to 20 ng/mL plasma).
- Vecuronium was diluted in PBSA to obtain a concentration of 0.18 μg/mL (corresponding to 0.30 μg/mL plasma).

One volume of the corresponding working solution was mixed with one volume of g and non-glycated blood samples, resulting in final concentrations of P at 4 μg/mL, R at 6 ng/mL (10 ng/mL plasma), and V at 0.09 μg/mL (0.15 μg/mL plasma)^26^. Table 9 summarizes the incubations performed with anesthetic agents. The samples were incubated at 37 °C for 30 minutes with slow, constant stirring. Due to R’s short lifetime, an additional dose was added after 15 minutes of incubation. Since RBCs do not metabolize P and V, no further doses of these anesthetics were required, maintaining their initial concentrations throughout incubation. Then, samples were centrifuged (1,500 g, 5 min, at room temperature) and washed three times with standard saline solution.

## Techniques and instruments

### Digital Analysis of Microscopic Images

An analysis of the morphology of erythrocyte aggregates (glycated and non-glycated) treated with anesthetics and controls was conducted. Images were captured using an Inverted Optical Microscope connected to a digital camera. The ISCapture software was employed for image processing. To capture the images, a 0.25% v/v suspension of red blood cells (RBCs) in autologous plasma was prepared, and five images were recorded in different fields for each sample. The following parameters were calculated for each image:

<u>Isolated Cell Coefficient (CCA)</u>: Defined as the difference between the percentage of individual cells before (CCA_initial_) and after treatment (CCA_final_) relative to the percentage of individual cells before treatment (CCA_initial_)^24^.

This coefficient can range from 0 to 1, where CCA = 0 indicates no difference in aggregation before and after treatment, and CCA = 1 indicates complete aggregation after treatment. This parameter allows for a quantification of the aggregation in the treated sample relative to the control.

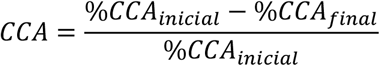

<u>Count and Classification of Erythrocyte Agregates</u>^22,23^: Aggregates were counted and classified into the following four categories:

% of individual cells (IC);

% of aggregates with 2, 3, or 4 cells;

% of aggregates with five or more cells;

% of large aggregate networks (AMAS acronym in Spanish).

The percentage of each category was calculated for each RBC suspension, and these percentages were averaged across all the studied samples.

## Aggregation Kinetics Parameters

This device, developed by the Biomedical Physics Group at IFIR (CONICET-UNR), obtained erythrocyte aggregation kinetics curves using the laser transmission technique. Characteristic curves of light intensity as a function of time during the aggregation kinetics process, known as silectograms, were generated^27,28^. Based on these curves, the following parameters were calculated for each sample:

Amp_t_ (amplitude): The amplitude of light intensity at a given time t, indicating the degree of red blood cell aggregation.

t_1/2_ (half time): The time required to reach half of the light intensity (Amp/2), indicating the characteristic time constant for an average level of aggregation.

Aggregation Index (AI): The ratio of the area under the silectogram to the total area over a specific period of time, indicating the normalized quantity of RBC aggregates.

For this assay, 15 µl of washed RBCs resuspended at 40% in autologous plasma were placed on the measurement slide, and the laser light intensity was recorded for 600 s.

## Stationary and Dynamic Viscoelastic Parameters

The Erythrocyte Rheometer^29–32^ was developed in the Laboratory of the Biomedical Physics Group at the Instituto de Física Rosario (IFIR - CONICET/UNR) and is based on the laser diffractometry technique. This instrument determined the stationary and dynamic (oscillatory) viscoelastic parameters of RBC membranes treated with anesthetics and controls (glycated and non-glycated). The geometric characteristics of the diffraction pattern are related to erythrocyte deformability and vary according to the applied shear stress. The loading and unloading curves were analyzed using specific software, which allows the calculation of the following parameters for each sample:

DI: deformability index

μ: elastic modulus of the RBC membrane η: surface viscosity of the RBC membrane

δ: phase shift between the applied shear stress and the erythrocyte response G**’**: dynamic elasticity or storage modulus

G**”**: dynamic loss modulus

η**’**: real component of complex viscosity

η**”**: imaginary component of complex viscosity

For these determinations, 110 µL of RBCs (washed and suspended at 40% in autologous plasma) were added to 4.5 mL of a 5% (w/v) polyvinylpyrrolidone (PVP) solution in PBS (viscosity: 22 ± 0.5 cP; pH: 7.4 ± 0.05; osmolality: 295 ± 8 mOsmol/kg; temperature: 25.0 ± 0.5 °C).

## Acknowledgements

The authors would like to extend their gratitude to the Universidad Nacional de Rosario and the Consejo Nacional de Investigaciones Científicas y Técnicas (CONICET, Argentina) for their support of this research. This work was specifically funded by the Universidad Nacional de Rosario, under project number BIO400 (Res. CS. N° 448/2017). Additionally, the first author, would like to acknowledge his doctoral scholarship from CONICET.

## Author contributions

MBS – conceptualization, methodology, investigation, formal analysis, original draft, and review & editing.

HVC - formal analysis, and review & editing.

NA - conceptualization, methodology, and review & editing.

BDR - conceptualization, methodology, review & editing, supervision, and funding acquisition AA - conceptualization, methodology, review & editing, supervision, and funding acquisition.

